# Multisensory integration across reference frames with additive feed-forward networks

**DOI:** 10.1101/2024.11.25.625309

**Authors:** Arefeh Farahmandi, Parisa Abedi Khoozani, Gunnar Blohm

## Abstract

The integration of multiple sensory inputs is essential for human perception and action in uncertain environments. This process includes reference frame transformations as different sensory signals are encoded in different coordinate systems. Studies have shown multisensory integration in humans is consistent with Bayesian optimal inference. However, neural mechanisms underlying this process are still debated. Different population coding models have been proposed to implement probabilistic inference. This includes a recent suggestion that explicit divisive normalization accounts for empirical principles of multisensory integration. However, whether and how divisive operations are implemented in the brain is not well understood. Indeed, all existing models suffer from the curse of dimensionality and thus fail to scale to real-world problems. Here, we propose an alternative model for multisensory integration that approximates Bayesian inference: a multilayer-feedforward neural network of multisensory integration (MSI) across different reference frames trained on the analytical Bayesian solution. This model displays all empirical principles of multisensory integration and produces similar behavior to that reported in Ventral intraparietal (VIP) neurons in the brain. The model achieved this without a neatly organized and regular connectivity structure between contributing neurons, such as required by explicit divisive normalization. Overall, we show that simple feedforward networks of purely additive units can approximate optimal inference across different reference frames through parallel computing principles. This suggests that it is not necessary for the brain to use explicit divisive normalization to achieve multisensory integration.

**Significance Statement:** This research presents an alternative model to divisive normalization models of multisensory integration in the brain. Our study demonstrates that a feed-forward neural network can achieve optimal multisensory integration across different reference frames without explicitly implementing divisive operations, challenging the long-held assumption that such operations are necessary for multisensory integration. The model displays all the empirical principles of multisensory integration, producing similar behavior to that reported in Ventral intraparietal (VIP) neurons in the brain. This work offers profound insights into the putative neural computations underlying multisensory processing.

## Introduction

Uncertainty is tightly linked to human sensory perception and interaction with the environment (Atkins et al., 2001). For example, when we are moving around on a foggy day, our visual information is less reliable compared to a sunny day. Uncertainties can arise from the information of different sensory modalities and the natural noisiness of spiking neurons (Faisal et al., 2008). Numerous behavioral studies showed that the brain integrates different sensory information in an optimal way to compensate for the uncertainties (Hillis et al., 2004). Multisensory integration in the nervous system is akin to a Bayesian inference process (Ernst and Banks, 2002; Ernst and Bülthoff, 2004; Knill and Pouget, 2004; Körding and Wolpert, 2006; Stein and Stanford, 2008; Battaglia et al., 2003; Merfeld et al., 1999; Wolpert et al., 1995; Meredith and Stein, 1986; Kojima and Landy, 2001; Landy et al., 1995; Kersten and Schrater, 2002). Since each sensory modality encodes information in a different coordinate system, reference frame transformations are necessary to achieve coherent multisensory integration (Deneve and Pouget, 2004). However, despite the abundant indications that the brain performs near-optimal probabilistic inferences, its neural implementation is a subject of debate.

Previous studies have proposed that the brain can perform statistical inference by marginalization over variables using explicit divisive normalization (Beck et al., 2011; Ohshiro et al., 2011; Pitkow and Angelaki, 2017). To elaborate more, optimal cue integration is considered an example of statistical inference in which different sensory signals are corrupted by uncertainty. Ohshiro et al. (2011) showed that a neural network implementing explicit divisive normalization at the multisensory level develops neuronal behavior reported in many brain areas involved in multisensory integration. Additionally, the authors took a further step and compared the prediction of a divisive normalization model with a model based on subtractive inhibition and showed that the divisive normalization provides a better prediction for the observed neuronal patterns (Ohshiro et al., 2017). Other neurophysiological studies, combined with computational modeling, provided evidence for the functional role of divisive normalization throughout the cortex, leading to the suggestion that explicit divisive normalization is a canonical cortex function (Carandini and Heeger, 2011).

Despite the established critical role of divisive normalization, how the brain implements such processes is a puzzle. This is mainly because explicit divisive normalization requires intractable division and multi-plication operations, making such implementation physiologically infeasible (Pitkow and Angelaki, 2017). Therefore, current models that utilize divisive normalization or, more broadly, statistical inferences, encounter various limitations. These limitations include a neat preconfigured connectivity structure (Deneve et al., 2001; Beck et al., 2011; Ohshiro et al., 2011), explicitly matching population codes for individual neurons (Ma et al., 2006; Ohshiro et al., 2011; Beck et al., 2012), similarity of all population codes (Ma et al., 2006), and/or the requirement of an unrealistically large number of neurons (Beck et al., 2011). Consequently, how divisive normalization can be implemented in a biologically feasible manner is unknown.

In this study, we chose a multisensory integration task across reference frames to investigate this issue. Specifically, the task was to estimate the position and the associated variability of the hand position across different eye positions using both retinal and proprioceptive sensory information. Previous networks required perfectly aligned and similar population codes to implement probabilistic inference, but we designed a task to merge retinal hand position with proprioceptive hand position with different neural coding schemes (i.e., probabilistic spatial and joint codes) within the same network. In addition, While divisive normalization is suggested for cue integration and coordinate transformations, its explicit form requires different neurons for each task (Emin Orhan and Ma, 2017). Here, we investigated if divisive normalization is inherently performed when all network units perform the same neuronal operations (Beck et al., 2012).

We trained a multi-layer feedforward neural network with standard error-based feedback to perform a multisensory integration task across reference frames. Not surprisingly, we show that this network can perform near-optimal probabilistic inference without requiring quadratic and divisive operations (Emin Orhan and Ma, 2017). We show that our network produces a wide range of behaviors similar to recorded neuronal activity in the brain. These behaviors include inverse effectiveness, the spatial correspondence principle, gain-like modulations, super-additivity, and multisensory suppression. In addition to key empirical principles of multisensory integration, it accounts for quantitative features of cue combination: we observed modulation of neural activity in our network by varying the cue reliability, similar to area MSTd (Morgan et al., 2008). The results of this study demonstrate that simple feed-forward networks of purely additive units can implement multisensory integration in the brain without the requirement of explicit divisive normalization.

## Materials and Methods

### Task

The goal of our model is to perform multisensory integration across reference frames. The task is to estimate the position and associated variability of the hand across different eye positions when visual and proprioceptive sensory information of hand position and proprioceptive information of eye position are available. Many previous studies showed that humans integrate information coming from different sensory modalities to decrease their uncertainty (Atkins et al., 2001; Ernst and Banks, 2002; Ernst and Bülthoff, 2004; Kersten et al., 2004; Knill and Pouget, 2004; Körding and Wolpert, 2004; Kojima and Landy, 2001; Landy et al., 1995; Stein and Stanford, 2008). Furthermore, varying eye orientation will bias retinal information demanding a reference frame transformation to compensate for eye orientation (Blohm and Crawford, 2007; Crawford et al., 2004). Therefore, to analytically combine visual and proprioceptive hand position, the retinal hand position is first updated based on the eye position and then optimally integrated with proprioception. The overall steps of our simple 1-dimensional proof-of-concept task are illustrated in Figure 1.C.

**Figure 1.**
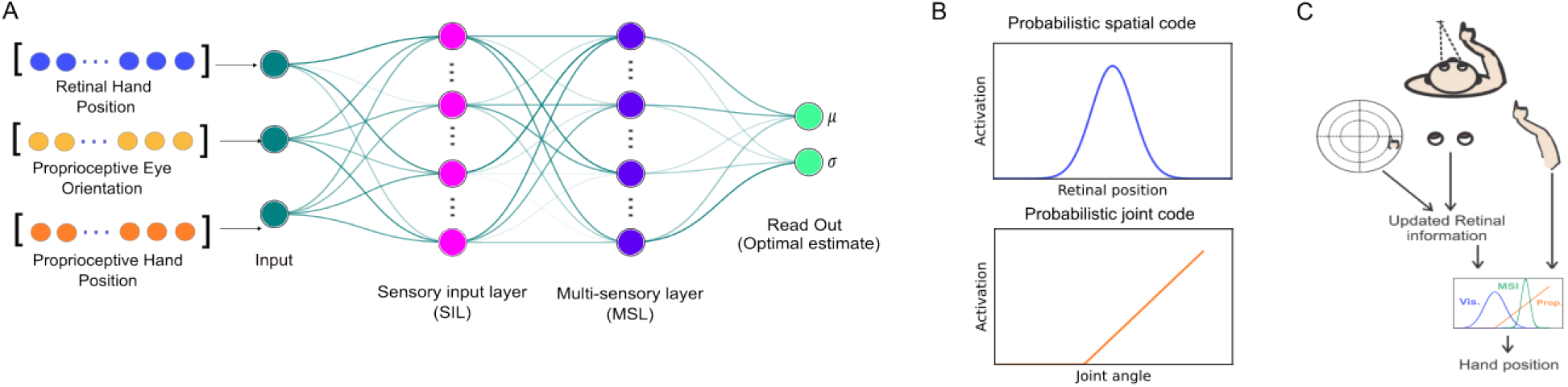
Network architecture and Task. **A**) The network consists of four layers: input layer, two hidden layers (sensory input layer (SIL) and multisensory layer (MSL), and a linear read-out layer. **B**) We used probabilistic spatial code to encode visual information and used probabilistic joint code to encode proprioceptive information of both hand position and eye orientation. **C**) The task is to estimate the hand position by combining the visual and proprioceptive sensory inputs across different eye orientation.

To generate the relevant input/output training data for the network, we assumed that all the representations have a Gaussian distribution with a mean and variance (visual hand position: 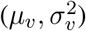, proprioceptive hand position: 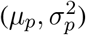, and eye position: 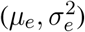 and are statistically independent. Therefore, the mean and variance of updated visual hand position 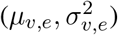 can be derived as following:

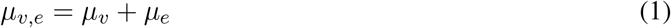

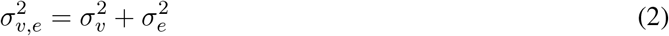

After transforming the retinal hand position into spatial hand position, the hand position is estimated by integrating the visual and proprioceptive information of hand position. Since both signals have Gaussian distributions and are independent, the mean and variance of the combined signal 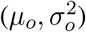 can be derived as follows (naive Bayesian integration):

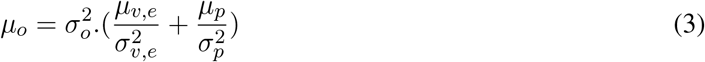

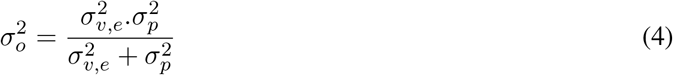

By substituting equations 1 and 2 into equations 3 and 4, we can find the relationship between estimated hand position (our network output) and sensory information (our network inputs):

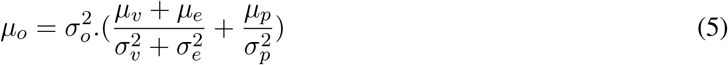

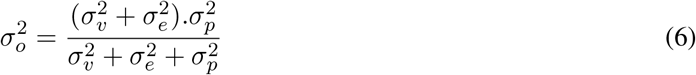

### Network Architecture

Our proposed network consisted of four layers: an input layer, two hidden layers, and a read-out layer (Figure 1.A). To generate the input layer, we had three groups of sensory input nodes: visual and proprioceptive information of hand position and proprioceptive information of eye position. The units in both input and read-out layers had linear transfer functions. There were two hidden layers in our network, each containing 64 units with sigmoid transfer function. The hidden layers included a sensory input layer (SIL) that all the sensory nodes projected to and a multisensory layer (MSL) which was designed to receive the input from the sensory input layer (SIL) and estimate the position of the hand using population coding. Then, the read-out layer mapped the population code to the desired mean and variance of the theoretical estimated hand position based on Bayesian integration (see above Task section). The network has both negative and positive weights between different layers. In the following, we provide technical details of the information coding.

### Input layer

We have three different sensory inputs for our network: visual and proprioceptive information of hand position as well as proprioceptive information of eye position. Therefore, our network consists of three groups of sensory input units coding for visual, proprioceptive, and eye positions. Visual information was encoded using visual-like tuning curves, while proprioceptive information of both hand and eye was coded using muscle-like tuning (Figure 1.B).

The visual information of hand position was encoded using a receptive field-like population coding scheme. We assumed that visual information is sampled from a normal distribution with mean *x*_*v*_ and variance 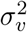. Each node in the visual group had a Gaussian tuning curve:

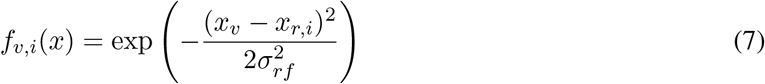

Where *f*_*v,i*_(*x*) is the firing rate of the *i*^*th*^ neuron in the visual sensory group, *x*_*r,i*_ is the neuron’s center of receptive field, and 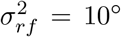 is the width of the receptive field. We assumed that the receptive field of all neurons in each sensory input group has the same width. Neurons were distributed uniformly along the visual field (−75°, 75°) and therefore the distance between neurons’ response field Δ*x* was derived as follows:

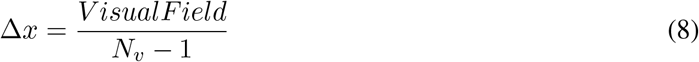

In which *N*_*v*_is the number of neurons in the visual sensory group. In addition, we assumed that the activation of each neuron was modulated based on the reliability of visual information and therefore the activity *a*_*i*_ of each neuron multiplied by a gain factor (in which *K* = 50 is a constant):

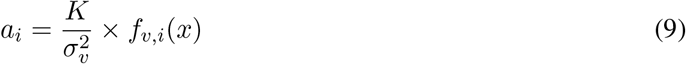

In addition, we used random Poisson noise to include biologically realistic trial-to-trial firing rate variability. Therefore, the final activation of each node was sampled from a Poisson distribution with *λ* = *a*_*i*_. Similar coding was used in previous neural network studies (Blohm et al., 2009; Zipser and Andersen, 1988; Scott Murdison et al., 2015).

During static position coding, tuning curves of neurons in motor cortex can be simplified to a linear, monotonic function (Paninski et al., 2004). Therefore, we used a simple linear tuning curve for coding both hand and eye positions. Furthermore, both positive and negative slopes were used for a unique mapping of position into neural coding based on push-pull mechanism (Blohm et al., 2009; Murdison et al., 2017; Xing and Andersen, 2000):

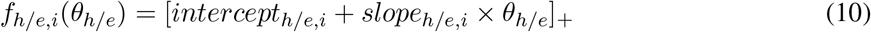

In which *f*_*h/e,i*_(*θ*_*h*/*e*_) is the firing rate of *i*^*t*^*h* neuron for proprioceptive hand (*h*) or eye (*e*) sensory input. Intercepts and slopes are chosen randomly between (−1,1) and (−10, 10) respectively for each unit. Joint angle and eye orientation then coded linearly in the activity of these units. Similar to visual information coding, we used reliability-based gain modulation (Eq. 9) and Poisson noise to generate the final neuronal activity of these units.

### Hidden layers

Our network contained two hidden layers: sensory input layer (SIL), which all the sensory units project to and multisensory layer (MSL), which receives the input from the previous layer and contains the estimated hand position in population codes. We used sigmoid transfer function for both hidden layers to mimic the non-linear transfer function of real neurons (Naka and Rushton, 1966). Specifically, we used the following equation to characterize the input (*x*) and output relationship *a*(*x*):

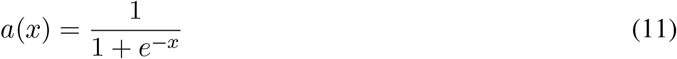

### Read out

As mentioned before, the goal of our network was to estimate hand position by integrating visual and proprioceptive information while accounting for different eye orientations. In the task section, we provided the theoretical relationship between input information and the estimated hand position 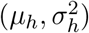 which is in accordance with behavioral (psychophysics) studies. Our network (presumably) generated the estimated hand position in population coding format at the MSL hidden layer. In order to translate population codes to their behavioral (psychophysics) counterpart, we added a linear layer to our network with two units, one for the mean value and one for the variance of the estimated hand position. The logic was that if the MSL represented correct estimates in a distributed fashion, then we should be able to read them out directly (like other brain areas would do). Therefore, our network automatically learns to map the population code to the desired mean and variance outputs.

### Data generation and network training

All the sensory input signals (visual, proprioceptive, and eye) were generated using random number generator in MATLAB 2022 (randn.m). Specifically, we used normal distributions N(0,15), N(0,18), N(0,20) for visual, proprioceptive, and eye positions respectively. Similarly, the visual, proprioceptive, and eye variance were randomly sampled from the range (1,40), (1,64), and (1,64) respectively. The output was generated based on the theoretical framework provided in the task section. We generated 50000 trials for training the network. A Resilient backpropagation method (Riedmiller and Braun, 1993) was used to train the network.

### Data Analysis

We performed several analyses to assess the extent to which our network model (and specifically our HLUs) replicated the reported neuronal activities associated with multisensory integration and reference frame transformations. These analyses were similar to those used in previous works (Avillac et al., 2007; Blohm et al., 2009; Morgan et al., 2008; Murdison et al., 2017; Ohshiro et al., 2011).

### Network performance

A quantitative analysis is performed to assess how well the network estimated the position of the hand and the associated uncertainty (variance). To do so, we simulated our network with 5000 randomly generated inputs that are not included in the training set. Then, we performed linear regression of read-out values (mean and variance of hand position) with the predicted values from our analytical models to examine how well our network is able to marginalize the eye orientation and estimate hand position.

### Multisensory integration empirical findings

In this work, we compared the activity of the units in the hidden layer of our network to the reported neuronal activity associated with multisensory integration. To examine to what extent our network’s implementation is comparable to an explicit divisive normalization network, we used the same parameters as previous studies (Avillac et al., 2007; Morgan et al., 2008; Ohshiro et al., 2011) to quantify the activity of hidden layer units: additivity index, response additivity, and response enhancement. These three parameters have been used in the literature to quantify the effect of multisensory integration at the neuronal level. All parameters quantify how bimodal activity is different compared to the activity when each stimulus is presented alone. We selected these parameters to be able to compare our results with the current literature.

1. Additivity index (AI): it compares how bimodal activity is enhanced compared to the sum of the activities to unimodal stimuli. A positive value of AI show to what extent the bimodal is enhanced and larger than the summation of unimodal responses. Conversely, if AI is negative, it showes how much the bimodal response is suppressed and less than sum of unimodal responses.

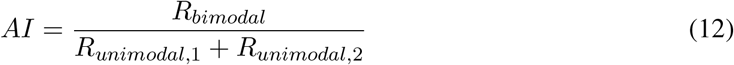
2. Response enhancement (RE) or Amplification index: it has been shown that generally neurons in the multisensory areas are most responsive to one of the stimuli (Avillac et al., 2007). This parameter compares how the response of a unit to a most effective stimulus is affected when both stimuli are presented. In other words, this parameter evaluates to what extent representing the non-effective stimulus suppresses or enhances the response of the effective stimulus. A positive value of this index represents multisensory enhancement, meaning that the second stimulus enhanced the response of the neuron. Also, a negative value represents multisensory suppression, meaning that the second stimulus suppressed the response of the neuron.

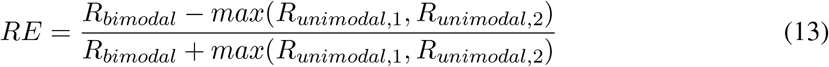
3. Response additivity (RA): this parameter compares the bimodal activity to the arithmetic sum of uni-modal activities. A positive value of response additivity represents superadditivity and a negative value represents sub-additivity. Superadditivity means the neuronal response is higher than the summation of responses to individual stimulus, and subadditivity means that the neuronal response is lower than the summation of responses to individual stimulus.

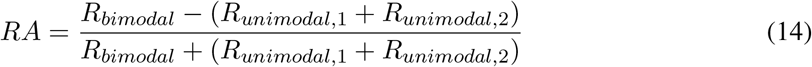

### Code Accessibility

Code is available on github (https://zenodo.org/records/14213670).

## Results

In this study, we propose that a feedforward neural network which is trained to perform multisensory integration across reference frames is functionally similar to an explicit divisive normalization. To support our hypothesis, we first examine if our network is capable of performing the multisensory integration task across reference frames. In the next step, we compare the activity of our network units in the hidden layers with the reported neuronal activities from multisensory areas in the brain. Specifically, we investigate if our network prediction is comparable with the prediction of an explicit divisive normalization model (Ohshiro et al., 2011). Finally, we explore the possible mechanisms that our network might use to perform multisensory integration.

### Network performance

First, we evaluated the performance of our network to confirm that our network has learned to perform the relevant aspect of the task before analyzing the activity of hidden units. To do so, we compared the predicted mean and variance of the hand position with the analytical solution of Bayesian integration (Figure 2). The regression analysis of the position revealed that the predicted values of our network match the desired value with good precision (Figure 2.A, slope 0.99 for the regression fit and *R*^2^ = 0.89). Additionally, we calculated the error in hand position estimation and observed that the mean error was relatively small (Figure 2.B : *µ* ≈ 7°, *σ* ≈ 6°). Similar results were observed for the variance of hand position estimation (Figure 2.C-D). These results indicate that our network accounted for the added uncertainty due to coordinate transformations and contributed this uncertainty into the integration process.

**Figure 2.**
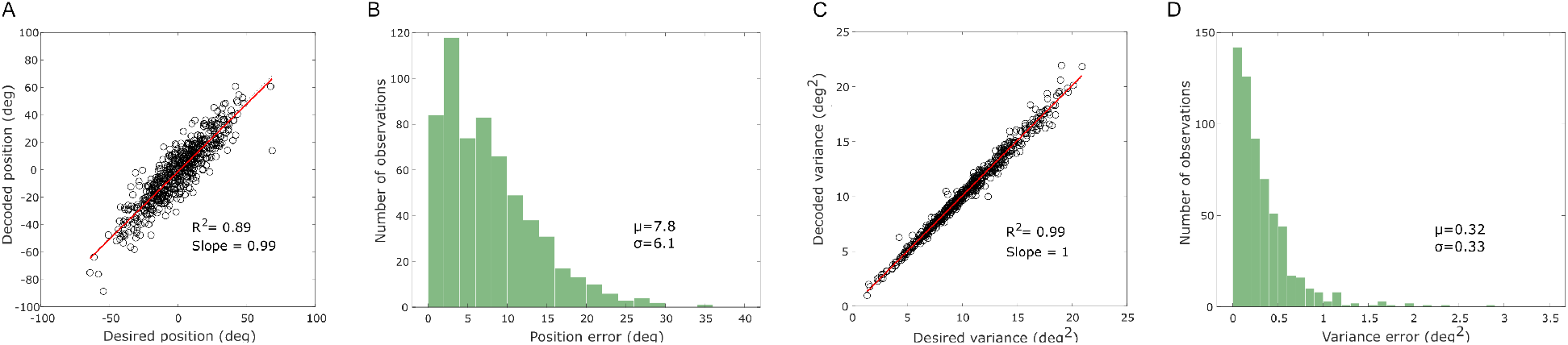
Network performance. **A**) Predicted hand position by the network compared to the desired position which is derived based on the analytical solution. **B**) Distribution of the position estimation errors produced by the network for a random set of test inputs (visual and proprioceptive hand positions and proprioceptive eye orientations) **C**) Predicted uncertainty is associated with hand position estimation compared to the desired value predicted by the analytical solution. **D**) Distribution of the uncertainty estimation errors produced by the network for a random set of test inputs.

After establishing that our network is able to perform multisensory integration, we aim to examine whether the units of the network mimic the neural activations reported in the brain regions involved in multisensory integrations (e.g. Ventral intraparietal (VIP), medial superior temporal (MSTd), etc.).

### Inverse effectiveness

Numerous studies showed that neurons in multisensory areas show enhanced responses to bimodal inputs (multisensory enhancement) vs. to unimodal inputs (Meredith and Stein, 1986; Stein and Stanford, 2008; Perrault et al., 2003; Stanford et al., 2005). In other words, the multisensory response is stronger than the response to each individual stimulus. However, this enhancement is not linear and follows the inverse effectiveness principle. Inverse effectiveness is a reported phenomenon in multisensory areas of the brain which indicates that multisensory enhancement becomes stronger when both stimuli are weak and decreases by increasing the input intensity (Stanford et al., 2005; Stein and Stanford, 2008). Ohshiro et al. (2011) quantified this phenomenon through their divisive normalization model. They calculated the additivity index by dividing the bimodal response to the arithmetic sum of unimodal responses and showed that additivity index is larger for weak bimodal inputs. This means that multisensory enhancement of weak inputs is higher than the sum of activities measured from single inputs (superadditivity). However, for strong inputs this enhancement disappears and activity is equal (additivity) or lower (subadditivity) than the arithmetic sum of activities produced by single inputs (Ohshiro et al., 2011).

To assess if the units of our network demonstrated similar behavior, we varied the intensity (i.e. reliability in this paper) of our visual and proprioceptive inputs by varying their variances (see ‘Materials and Methods’ for further detail). Similar to Ohshiro et al. (2011), we calculated the additivity index for each unit in the multisensory layer by dividing the multisensory response to the sum of the two unimodal responses and showed the result for example units in Figure 3. In agreement with the divisive normalization model (Ohshiro et al., 2011), as the reliability of the unimodal inputs increases, the additivity index becomes weaker showing less multisensory enhancement for more reliable inputs. This inverse effectiveness phenomenon can be seen in Figure 3 in units with different bimodal response behaviour.

**Figure 3.**
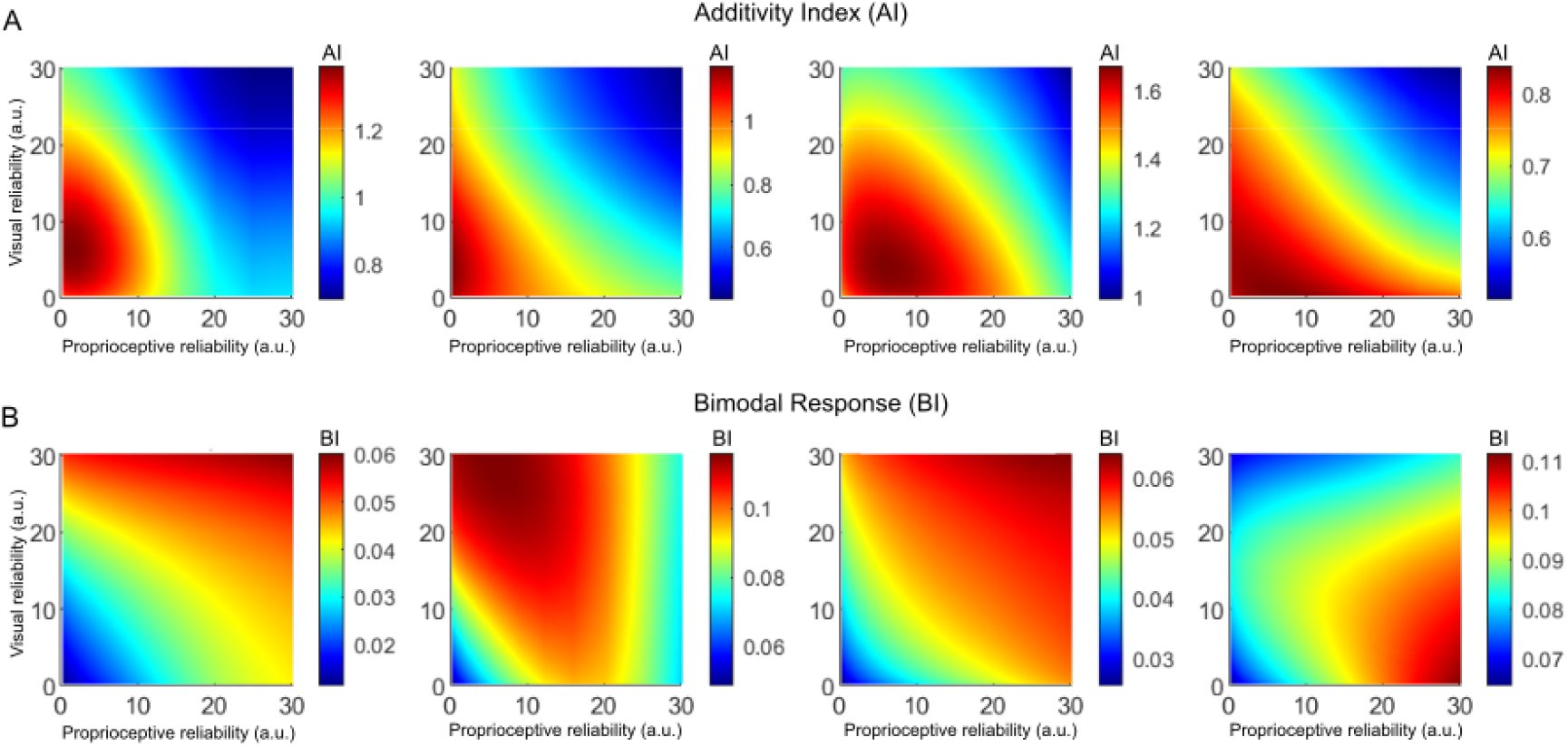
Inverse effectiveness in multisensory layer (MSL). **A**) Additivity index for some sample units in multisensory layer (MSL). **B**) Corresponding bimodal response of sample units with different reliability of input.

Our model only displays super-additivity (*AI* > 1) when the response to both unimodal stimuli is weak (Figure 3.A). As the intensity of the response to unimodal stimuli increases the multisensory response becomes sub-additive (or additive). Therefore, the multisensory layer of our network generated behavior that was in agreement with findings reported from multisensory areas of the brain such as the superior colliculus (Alvarado et al., 2007; Perrault et al., 2005). Note that this is only true for the Multisensory layer of our network; the units in the Sensory-input layer of our model are not producing inverse effectiveness. Therefore, similar to the explicit divisive normalization model (Ohshiro et al., 2011) our network reveals inverse effectiveness regardless of whether weak inputs produce superadditivity or not, as shown in Figure 3.B with different bimodal response behaviours (Stanford et al., 2005; Perrault et al., 2005).

In addition, we used a complementary way, similar to Avillac et al. (2007), to further determine the inverse effectiveness. Avillac et al. (2007) calculated an amplification index which is the percentage of response enhancement for integrative neurons in the VIP region. They showed this index as a function of the dominant unimodal response with the same input for units with enhanced multisensory responses to assess inverse effectiveness (Figure 4.B). They observed that as the dominant unimodal response increases, the amplification index decreases as an indicator of the inverse effectiveness principle. We calculated the same index for all enhanced units in the multisensory layer with different inputs and plotted them against the dominant unimodal response. Figure 4.A shows a similar observation of inverse effectiveness in our model; the amplification index decreases as the dominant unimodal responses decrease.

**Figure 4.**
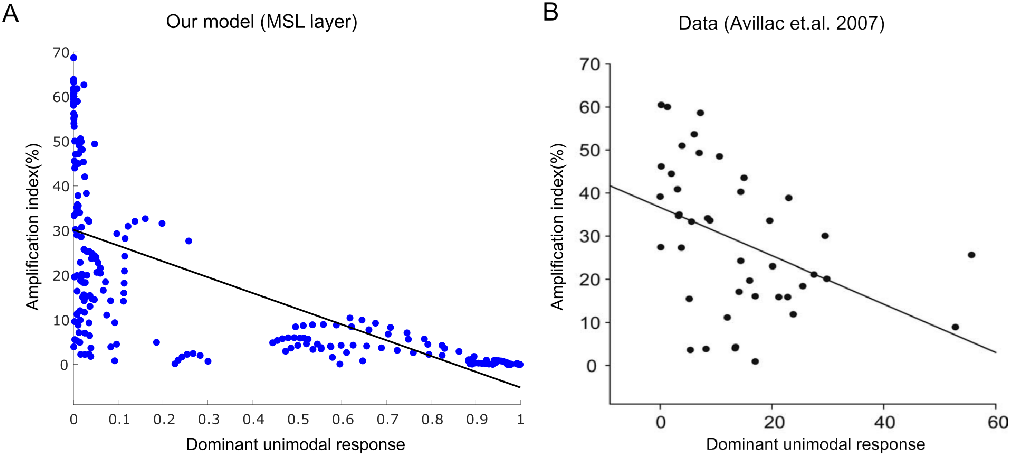
Inverse effectiveness principle in our model and data (VIP). The amplification index is plotted as a function of the dominant unimodal response for the subset of the integrative units in **A**) our model and **B**) area VIP with enhancive multisensory responses. (replotted with permission from Avillac et al. (2007))

### Cross-modal enhancement and suppression in bimodal and unimodal responses

Neurophysiological recordings in multisensory regions (such as the superior colliculus (SC) and the ventral interparietal area (VIP)) illustrated that neurons show both multisensory enhancement and suppression (Avillac et al., 2007; Meredith and Stein, 1986). Among these studies, Avillac et al. (2007) quantified neuronal behavior in area VIP of monkeys using both response enhancement and response additivity (see ‘Materials and Methods’ for further details). Using these two parameters, Avillac et al. (2007) showed that VIP neurons act heterogeneously, exhibiting both cross-modal enhancement and suppression along with nonlinear super- and sub-additivity (Figure 5.C). To examine whether our model replicates the reported behaviors, we simulated the trained network to perform multisensory integration while varying the retinal and proprioception information of the hand for a fixed position and reliability of the eyes. We selected 11 uniformly distributed positions for both the retinal and proprioceptive positions in the range of trained positions. Then, we calculated both response enhancement and response additivity for all the possible combinations of the different retinal and proprioceptive information of the hand position as shown in Figure 5.A-B for both the MSL and SIL layers. As Figure 5.A illustrates the units in the SIL layer only show cross-modal suppression and sub-additivity. However, the units in the MSL layer of our network replicated a similar variability as reported in VIP neurons (Figure 5.B). For further investigation of units in the MSL layer, the enhancement and additivity of bimodal responses is plotted as a function of the reliability of the retinal and proprioceptive hand information. We show that the average of both the enhancement and additivity index decreases for more reliable sensory information, which is also consistent with inverse effectiveness (Figure 5.D).

**Figure 5.**
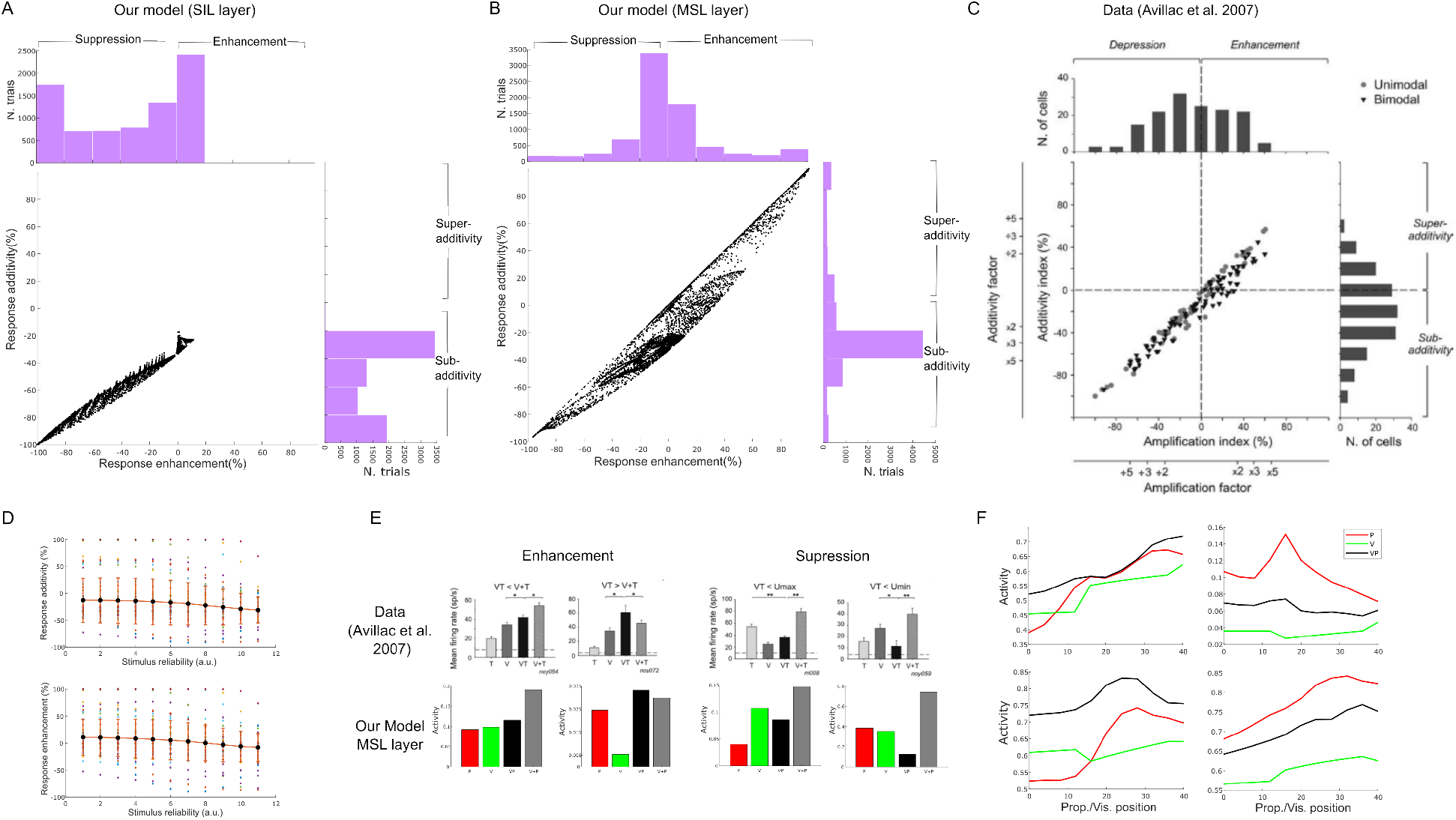
Multisensory enhancement and suppression in our model. **A**) Units in the SIL layer of our network only showed cross-modal suppression. **B**) Units in the MSL layer showed behavior comparable to recordings from area VIP and prediction of divisive normalization. **C**) Summary of the reported data from area VIP (Avillac et al. (2007)). **D**) Response enhancement and additivity are plotted as a function of stimulus reliability. The average of both indices decrease by increasing relaiability of stimulus in units of MSL layer. **E**) A comparison between sample units of our MSL layer and neurons in area VIP showing different responsiveness to the visual stimulus. Representing the proprioceptive stimulus alongside with the visual stimulus resulted in suppressed bimodal activity (i.e. cross-modal suppression) or Cross-modal enhancement. V, P: visual and proprioceptive unimodal responses; VP: Bimodal response; V+P: summation of unimodal responses. **F**) Recording from sample units of our MSL layer revealed the existence of units responsive to both stimuli or only to one of the stimuli (i.e. bimodal and unimodal neurons).

In addition to the observed population heterogeneity, Avillac et al. (2007) observed that neurons in area VIP can be categorized as bimodal or unimodal neurons. That is, bimodal neurons are determined as neurons that are responsive to both stimuli when presented alone. However, unimodal neurons respond only to one of the stimuli. Interestingly, unimodal neurons showed multisensory enhancement and suppression, similar to bimodal neurons. As shown in Figure 5.B, the MSL of our network replicated the population behavior of VIP neurons. However, that population behavior could result from merely bimodal neurons. Therefore, we analyzed individual units in the MSL of our simulated network with the aim of unraveling the details of each unit’s behavior (Figure 5.F). Figure 5.F illustrates, the MSL layer of our network contains some units behaving similar to unimodal neurons as well as other units behaving like bimodal neurons. For example, Figure 5.F top right shows a unimodal unit and Figure 5.F top left shows a bimodal unit in the MSL layer. Also, unimodal units exhibit both cross-modal suppression and cross-modal enhancement (Figure 5.E). In general, the MSL units of our network replicate both population and individual neuronal activity reported in the area VIP (Avillac et al., 2007).

### Multisensory integration and cue reliability

Extensive work has shown that multisensory integration at the behavioral level can be explained by a Bayesian framework (Ernst and Banks, 2002). In special cases, when both modalities have Gaussian distributions and are independent, the cue combination can be modeled as a weighted sum of the two stimuli and the weights are determined based on stimulus reliability (Atkins et al., 2001; Ernst and Banks, 2002; Ernst and Bülthoff, 2004; Kersten et al., 2004; Knill and Pouget, 2004; Körding and Wolpert, 2004; Landy et al., 1995; Stein and Stanford, 2008; Kojima and Landy, 2001). That is, the stimulus with higher reliability would have a higher weight (equivalent to a higher contribution) in the integration. Interestingly, Morgan et al. (2008) established that multisensory integration in the macaque visual cortex depends on stimuli reliability and showed that modulating cue reliability changes the activity of multisensory neurons in area MSTd. More specifically, the authors showed that neuronal responses to bimodal stimuli can be well explained by a weighted sum of neuronal responses to unimodal stimuli and that the weights are determined based on cue reliability: lower reliability of one of the stimuli results in a lower weight in integration (Morgan et al., 2008).

To examine if such a rule also holds among individual units in our network, we simulated the trained network varying one sensory stimulus reliability (here visual) while keeping the other sensory reliability fixed. We fixed eye position for this simulation and selected four levels of reliabilities: from low to high. We fixed the proprioceptive reliability at the medium level. Morgan et al. (2008) showed that the activity of neurons in area MSTd changes by decreasing the visual coherence (decreasing reliability). Figure 6.A-B shows the receptive field of a sample unit from the SIL layer of our network for a low and a high visual reliability, respectively. As can be seen, the neuronal response of this sample unit changed due to changes of the reliability of visual information. A sample unit in the MSL of our network displayed a similar behavior (Figure 6.C-D). In other words, the activity of units in our network is modulated by varying the reliability of visual information. This is an evidence that different sensory inputs are combined through gain modulation to perform reference frame transformation in multisensory integration tasks (Blohm and Crawford, 2009; Blohm, 2012). To quantify cue reliability at the population level, we fitted the bimodal response of each network unit with the weighted sum of the two unimodal responses:

**Figure 6.**
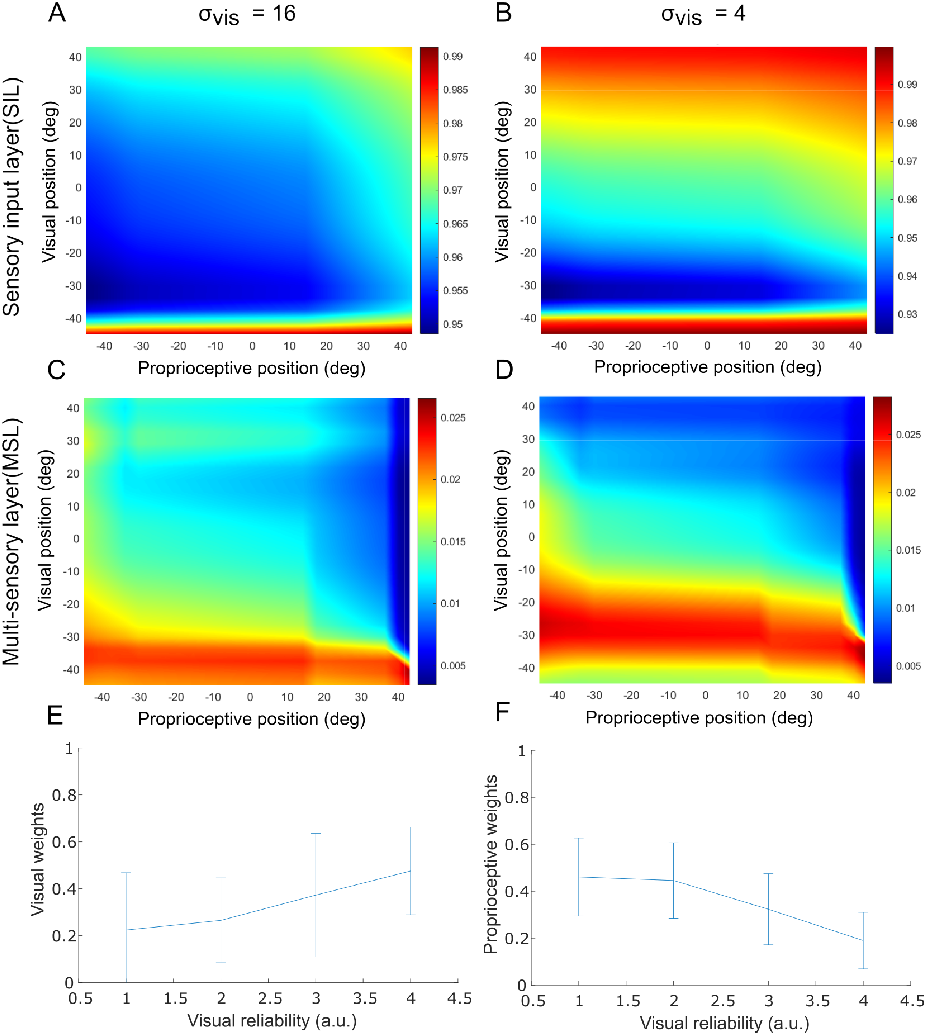
Cue reliability changes the response of multisensory integration. **A-D**) Receptive field for two sample units in the SIL (A-B) and MSL layer (C-D) for two different visual variabilities. The activity of units in both layers is modulated by varying the reliability of visual information. **E-F**) The increase of visual reliability resulted in an increase in the visual weights (E) and a decrease in proprioceptive weights (F).

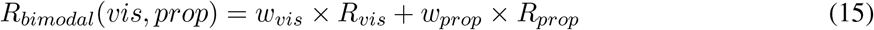

The visual and proprioceptive weights were calculated for each of the visual reliabilities. The linear fit was a good approximation to responses of the network units, with average *R*^2^ values of 0.68, 0.67, 0.69 and 0.70 for the 4 simulated reliabilities respectively. Similar to Morgan et al. (2008), increasing the visual reliability resulted in increased visual weights (Figure 6.E) and analogously decreased proprioceptive weights (Figure 6.F). These results are comparable to reported data from area MSTd and the explicit divisive normalization model predictions (Ohshiro et al., 2011).

### Gain modulation in multisensory integration

Previous studies have suggested that gain modulation could serve as a general purpose mechanism to implement multisensory integration across different reference frames (Andersen et al., 1985; Blohm et al., 2009; Blohm and Crawford, 2009; Blohm, 2012; Chang et al., 2009; Deneve and Pouget, 2004; Ma et al., 2006; Salinas and Sejnowski, 2001; Zipser and Andersen, 1988; Scott Murdison et al., 2015). Gain fields are defined as a multiplicative factor that scales the receptive field of a neuron up / down (Blohm and Crawford, 2009). As we saw cue reliability in units of our network, we can evaluate gain modulation and compute gain indices for our units. To do so, we calculated the receptive field (RF) of each unit, similar to Figure 6.A-D, by varying the retinal and proprioceptive hand positions for different eye rotations along with the associated variabilities. We observed that our units in both the SIL and the MSL are modulated by different visual or proprioceptive positions, thus mimicking the receptive field-like behavior (Figure 7.A-J upper panels). In the next step, we computed gain modulation index across the whole receptive field (Blohm, 2012):

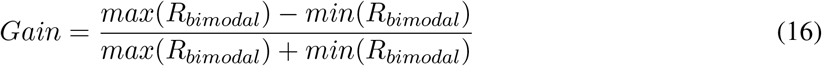

lower panels in Figure 7.A-J shows the distribution of gain indices for all units by varying different parameters such as eye orientation in both the SIL and MSL layers. this observed gain modulation could potentially explain the functionality of our network in performing multisensory integration across different reference frames.

**Figure 7.**
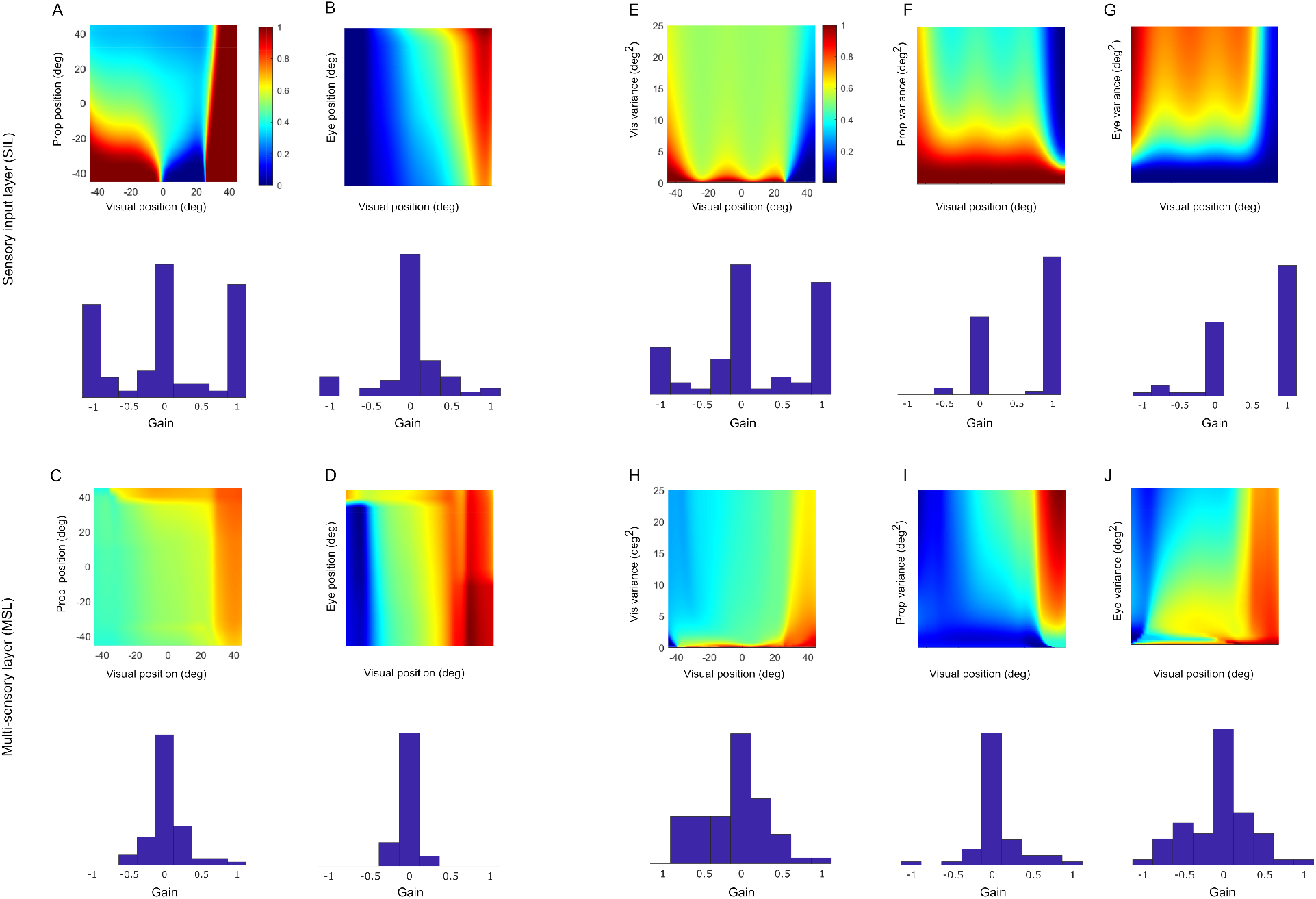
Gain Modulation. Gain modulation is characterized as up-/down-regulation of RFs by a secondary input without shifting the location of the RF. Upper panels represents unit example while lower panel represent population response. Units displayed gain modulation in both layers of the network. Additionally, the units in MSL showed a combination of shifted RF and gain modulations. **A-D** varying proprioceptive and eye position information caused gain modulation of the visual RF of the units in both the SIL and the MSL. **E-J** varying the variability of visual, proprioceptive, and eye-orientation information caused gain modulation of visual RFs of units throughout the network.

## Discussion

We proposed a distributed neuronal computation model to perform statistical inference. As a proof of concept, we trained a feedforward artificial neural network to implement multisensory integration across different coordinate frames and showed that the network is capable of performing the inference task with a small number of units without a neatly organized and regular connectivity structure required by explicit divisive normalization. Furthermore, analyzing the emergent properties of our network showed similarities with reported neurophysiological behaviors of neurons in multisensory areas in the brain, such as SC, MSTd and VIP. Specifically, we showed that our network reproduced inverse effectiveness, cross-modal suppression, and multisensory suppression of unimodal neurons. In addition, we showed that the activity of units in our trained network is modulated by cue reliability. This provides an interpretable explanation for multisensory integration across reference frames.

### Critical comparison with other models

Much previous work has focused on multisensory integration and coordinate transformation as an example of statistical inference (Deneve et al., 2001; Deneve and Pouget, 2004; Ma et al., 2006; Beck et al., 2011). This line of studies suggests that a population of neurons represent a probability distribution of the stimuli rather than a deterministic value. This coding is called probabilistic population coding (PPC) (Beck et al., 2011). The authors then show that a particular class of neural networks, basis function networks with multidimensional attractors, can implement probabilistic inference for many tasks. Additionally, in their later work, the authors established that for a specific class of neuronal variability, i.e. independent Poisson noise, there is a close-form solution for tasks such as cue integration (Ma et al., 2006; Deneve et al., 2001). That is, cue integration can be implemented in a simple neuron by neuron summation of sensory inputs. Similar to this work, we also deployed PPCs to code sensory information. Namely, we assumed that populations of neurons code the probability distribution rather than single values. However, we did not consider the same representation of population coding for all inputs and outputs. We allowed our network to learn the feedforward weights and consequently its own basis function. This implementation resulted in fundamental differences between our model and basis function networks. The first key distinction is that our network can perform the task with significantly fewer units and connections resulting in lower computational complexity.

For instance, the proposed network by Deneve et al. (2001); Beck et al. (2011) requires *N*^*d*.*s*^ 10^12^ units for combining *s* = 2 cues in 3-D space (*d* = 3, assuming each dimension is spanned by *N* = 100 units. In contrast, our network requires only *N*.*d*.*s* ≈ 10^3^ units for performing the same task, albeit with slightly lower accuracy. Furthermore, unlike basis function networks, our network supports different population coding representations for sensory inputs and outputs (Deneve et al., 2001; Beck et al., 2011). Finally, while basis function networks (Deneve et al., 2001; Deneve and Pouget, 2004; Pouget and Snyder, 2000) depend on strictly constrained connectivity structures between neurons, our network operates effectively without such restrictions.

As an alternative class of models for multisensory integration, explicit divisive normalization networks have been suggested (Ohshiro et al., 2011). One of the main strengths of such models is the ability to replicate many reported empirical principles of multisensory integration, including inverse effectiveness and cross-modal suppression, at the neuronal level. This is not the case for the probabilistic population code models discussed above (e.g. Ma et al. (2006)). However, these divisive normalization models, similar to PPC models, require neat connectivity configurations and a substantial number of synaptic connections for the normalization pool. More generally, explicit implementation of divisive normalization can result in intractable computations for which synaptic connectivity would virtually be infinite (Pitkow and Angelaki, 2017). Similar to explicit divisive normalization, our model reproduces reported multisensory empirical principles but without any explicit divisive operations. This led to a considerably lower complexity (i.e. required units and connections) of our proposed model compared to previous models.

A third category of multisensory integration models use dynamic neural networks with lateral inhibitions, subtractive normalization, and neuronal sigmoid transfer functions to reproduce both inverse effectiveness and cross modal suppression (Magosso et al., 2008; Ursino et al., 2009). It can be shown that, in such networks, inverse effectiveness mostly resulted from the sigmoid feature of neurons while cross modal suppression resulted from the lateral inhibition in the multisensory layer. Our network is similar to these networks in containing units with sigmoid transfer function; however, our network contains only feed-forward connections and no lateral connections. Nevertheless, our network is capable to replicate the reported cross-modal suppression through a much simpler, feed-forward process.

### Model limitations and potential future work

Our model has several limitations that can be the subject of future work. First, we used a very simple network: all units have the same sigmoid transfer function, and the network contains only feedforward connections. While the cortex has heterogeneity both with regard to connections (for example, feedforward, feedback, and lateral connections) and to cell types (Burkhalter and Bernardo, 1989; Darmanis et al., 2015; Harris and Shepherd, 2015). Future extensions of the model might shed light on the functional role of this heterogeneity. For instance, lateral connections can be added in different hidden layers of the network at the training stage. Then, by removing the lateral connections, we can analyze the functional role of the lateral inhibitions in different layers for statistical inference. In addition, this method could be used to investigate to what extent observed nonlinearities such as cross-modal suppression are dependent on lateral connections.

Another simplification of our model is that we only considered a specific class of coding: probabilistic populations codes (PPC) in which activity scales with reliability. That is, higher sensory reliability results in higher neuronal activity. Note that we define reliability as the inverse of variance of sensory inputs to be comparable with experimental studies. This definition is consistent with previous studies (Fetsch et al., 2011). Additionally, the neuronal variability was assumed to be Poisson. While this class of population coding is reported in many brain areas (Salinas and Abbott, 1994; Sanger, 1996; Seung and Sompolinsky, 1993; Snippe, 1996; Wu et al., 2001; Zemel et al., 1998), other classes of population coding have also been suggested (Fetsch et al., 2011; Krekelberg et al., 2006; Morgan et al., 2008). In a similar study, Emin Orhan and Ma (2017) implemented several statistical inferences with generative neural networks and showed that their network is able to perform the inferences without any constraint on the type of coding mechanisms used for sensory information. In the same way, we expect that changing the population coding should not affect the network’s ability to perform the task. However, it is not clear how different coding schemes for neural network inputs and outputs influence the specific solutions that a network generates to perform the multisensory integration task. This can be the topic of future investigations.

Lastly, we trained our network using back-propagation methods. There is a long-lasting debate in the literature on the feasibility of physiological systems to implement back-propagation (Crick, 1989; Grossberg, 1987), mainly due to its nonlocalality characteristics. However, recent work provided evidence that back-propagation can be estimated using more physiologically feasible functions (Bengio et al., 2015; Lillicrap et al., 2016). It would be valuable to investigate how much the results obtained depend on the exact training algorithm. However, we do not expect fundamental differences in the network’s performance, as previous studies have shown that different optimization methods tend to converge to similar solutions when reaching the global minimum (Blohm et al., 2009).

### Model implications

Our network replicates the empirical principles of multisensory integration exhibited by single neurons and is comparable to a neural network implementing explicit divisive normalization (Ohshiro et al., 2011). This implies that divisive normalization can be seen as an emergent functional characteristic of a system performing statistical inferences with noisy inputs in complex environments and is not necessary to implement statistical inference. One of the main aspects of inferences in such complex environments is marginalizing out the nuisance parameters. For example, in our model, eye orientation does not directly affect the cue combination; however, failing to account for eye orientation results in mis-estimation of the position of the hand. This observation might provide an explanation for the discrepancies between the PPC framework (Ma et al., 2006) and the explicit divisive normalization model (Ohshiro et al., 2011). Specifically, the PPC framework states that cue integration can be implemented by summation of individual neuron activity coding sensory information. However, this linear summation fails to explain the empirical principles of multisensory integration (i.e. non-linearities) at the neuronal level. On the other hand, explicit divisive normalization at the multisensory level can successfully replicate the empirical principles at the single neuron level. Based on our model, the observed non-linearities are emergent properties of the marginalization operations that are crucial for cue integration.

Our network implicitly implements divisive normalization with purely additive computations and in a feedforward manner. It does so by adapting the feed-forward weights during training (Chalk et al., 2018). This resulted in fixed decoding (read-out layer) and context-dependent dynamic receptive fields (RFs) in our network (gain modulation in Figure 7 and cue reliability in Figure 6). Intriguingly, such dynamic RFs have been reported in different areas of the brain (Fournier et al., 2011; Kabara and Bonds, 2001; Trott and Born, 2015; Yeh et al., 2009). These observations imply that marginalization through implicit divisive normalization might be a neuronal mechanism to adapt the activity in a way that the population code can be decoded by linear and fixed decoders, e.g. by higher cortical regions. This could explain the prevalence of apparent divisive normalization in many areas of the brain, such as the primary visual cortex, the olfactory system, and the hippocampus, and supports the speculation that it acts as a canonical operation throughout the cortex (Carandini and Heeger, 2011).

## Conclusion

The results of this study reveal that multisensory integration can be achieved in simple feedforward networks of purely additive units without the requirement of explicit divisive normalization. This is in line with the current view that the observed neuronal activities can be explained as emergent properties of the optimization processes required for statistical inferences.

## Acknowledgements

This work was supported by the Natural Sciences and Engineering Research Council of Canada and the Canada Foundation for Innovation.

## References

Alvarado JC, Vaughan JW, Stanford TR, Stein BE (2007) Multisensory versus unisensory integration: Contrasting modes in the superior colliculus. Journal of Neurophysiology 97:3193–3205.

Andersen RA, Essick GK, Siegel RM (1985) Encoding of spatial location by posterior parietal neurons. Science (New York, N.Y.) 230:456–458.

Atkins JE, Fiser J, Jacobs RA (2001) Experience-dependent visual cue integration based on consistencies between visual and haptic percepts. Vision Research 41:449–461.

Avillac M, Hamed SB, Duhamel JR (2007) Multisensory integration in the ventral intraparietal area of the macaque monkey. Journal of Neuroscience 27:1922–1932.

Battaglia PW, Jacobs RA, Aslin RN (2003) Bayesian integration of visual and auditory signals for spatial localization. Journal of the Optical Society of America. A, Optics, image science, and vision 20:1391.

Beck JM, Latham PE, Pouget A (2011) Marginalization in neural circuits with divisive normalization. Journal of Neuroscience 31:15310–15319.

Beck JM, Ma WJ, Pitkow X, Latham PE, Pouget A (2012) Not Noisy, Just Wrong: The Role of Suboptimal Inference in Behavioral Variability. Neuron 74:30–39.

Bengio Y, Lee DH, Bornschein J, Lin Z (2015) Towards Biologically Plausible Deep Learning. arXiv.org.

Blohm G (2012) Simulating the Cortical 3D Visuomotor Transformation of Reach Depth. PLOS ONE 7:e41241.

Blohm G, Crawford JD (2007) Computations for geometrically accurate visually guided reaching in 3-D space. Journal of Vision 7.

Blohm G, Crawford JD (2009) Fields of Gain in the Brain.

Blohm G, Keith GP, Crawford JD (2009) Decoding the cortical transformations for visually guided reaching in 3D space. Cerebral Cortex 19:1372–1393.

Burkhalter A, Bernardo KL (1989) Organization of corticocortical connections in human visual cortex. Proceedings of the National Academy of Sciences of the United States of America 86:1071–1075.

Carandini M, Heeger DJ (2011) Normalization as a canonical neural computation. Nature Reviews Neuro-science 13:51–62.

Chalk M, Marre O, Tkačik G (2018) Toward a unified theory of efficient, predictive, and sparse coding. Proceedings of the National Academy of Sciences of the United States of America 115:186–191.

Chang SW, Papadimitriou C, Snyder LH (2009) Using a Compound Gain Field to Compute a Reach Plan. Neuron 64:744–755.

Crawford JD, Medendorp WP, Marotta JJ (2004) Spatial transformations for eye-hand coordination. Journal of neurophysiology 92:10–19.

Crick F (1989) The recent excitement about neural networks. Nature 337:129–132.

Darmanis S, Sloan SA, Zhang Y, Enge M, Caneda C, Shuer LM, Gephart MG, Barres BA, Quake SR (2015) A survey of human brain transcriptome diversity at the single cell level. Proceedings of the National Academy of Sciences of the United States of America 112:7285–7290.

Deneve S, Latham PE, Pouget A (2001) Efficient computation and cue integration with noisy population codes. Nature Neuroscience 2001 4:8 4:826–831.

Deneve S, Pouget A (2004) Bayesian multisensory integration and cross-modal spatial links. Journal of Physiology Paris 98:249–258.

Emin Orhan A, Ma WJ (2017) Efficient probabilistic inference in generic neural networks trained with non-probabilistic feedback. Nature communications 8.

Ernst MO, Banks MS (2002) Humans integrate visual and haptic information in a statistically optimal fashion. Nature 2002 415:6870 415:429–433.

Ernst MO, Bülthoff HH (2004) Merging the senses into a robust percept. Trends in Cognitive Sciences 8:162–169.

Faisal AA, Selen LP, Wolpert DM (2008) Noise in the nervous system. Nature Reviews Neuroscience 2008 9:4 9:292–303.

Fetsch CR, Pouget A, Deangelis GC, Angelaki DE (2011) Neural correlates of reliability-based cue weighting during multisensory integration. Nature Neuroscience 2011 15:1 15:146–154.

Fournier J, Monier C, Pananceau M, Frégnac Y (2011) Adaptation of the simple or complex nature of V1 receptive fields to visual statistics. Nature Neuroscience 2011 14:8 14:1053–1060.

Grossberg S (1987) Competitive Learning: From Interactive Activation to Adaptive Resonance. Cognitive Science 11:23–63.

Harris KD, Shepherd GM (2015) The neocortical circuit: themes and variations. Nature Neuroscience 2015 18:2 18:170–181.

Hillis JM, Watt SJ, Landy MS, Banks MS (2004) Slant from texture and disparity cues: optimal cue combination. Journal of vision 4:967–992.

Kabara JF, Bonds AB (2001) Modification of response functions of cat visual cortical cells by spatially congruent perturbing stimuli. Journal of Neurophysiology 86:2703–2714.

Kersten D, Mamassian P, Yuille A (2004) Object perception as Bayesian inference. Annual Review of Psychology 55:271–304.

Kersten D, Schrater P (2002) Pattern inference theory: A probabilistic approach to vision na.

Knill DC, Pouget A (2004) The Bayesian brain: The role of uncertainty in neural coding and computation. Trends in Neurosciences 27:712–719.

Kojima H, Landy MS (2001) Ideal cue combination for localizing texture-defined edges. JOSA A, Vol. 18, Issue 9, pp. 2307-2320 18:2307–2320.

Körding KP, Wolpert DM (2004) Bayesian integration in sensorimotor learning. Nature 427:244–247.

Körding KP, Wolpert DM (2006) Bayesian decision theory in sensorimotor control. Trends in cognitive sciences 10:319–326.

Krekelberg B, Van Wezel RJ, Albright TD (2006) Interactions between Speed and Contrast Tuning in the Middle Temporal Area: Implications for the Neural Code for Speed. Journal of Neuroscience 26:8988–8998.

Landy MS, Maloney LT, Johnston EB, Young M (1995) Measurement and modeling of depth cue combination: in defense of weak fusion. Vision research 35:389–412.

Lillicrap TP, Cownden D, Tweed DB, Akerman CJ (2016) Random synaptic feedback weights support error backpropagation for deep learning. Nature communications 7.

Ma WJ, Beck JM, Latham PE, Pouget A (2006) Bayesian inference with probabilistic population codes. Nature Neuroscience 9:1432–1438.

Magosso E, Cuppini C, Serino A, Di Pellegrino G, Ursino M (2008) A theoretical study of multisensory integration in the superior colliculus by a neural network model. Neural Networks 21:817–829.

Meredith MA, Stein BE (1986) Visual, auditory, and somatosensory convergence on cells in superior colliculus results in multisensory integration. 10.1152/jn.1986.56.3.640 56:640–662.

Merfeld DM, Zupan L, Peterka RJ (1999) Humans use internal models to estimate gravity and linear acceleration. Nature 398:615–618.

Morgan ML, DeAngelis GC, Angelaki DE (2008) Multisensory Integration in Macaque Visual Cortex Depends on Cue Reliability. Neuron 59:662–673.

Murdison TS, Blohm G, Bremmer F (2017) Predictive orientation remapping maintains a stable retinal percept. bioRxiv p. 193250.

Naka KI, Rushton WA (1966) An attempt to analyse colour reception by electrophysiology. The Journal of Physiology 185:556.

Ohshiro T, Angelaki DE, Deangelis GC (2011) A normalization model of multisensory integration. Nature Neuroscience 14:775–782.

Ohshiro T, Angelaki DE, DeAngelis GC (2017) A Neural Signature of Divisive Normalization at the Level of Multisensory Integration in Primate Cortex. Neuron 95:399–411.

Paninski L, Fellows MR, Hatsopoulos NG, Donoghue JP (2004) Spatiotemporal Tuning of Motor Cortical Neurons for Hand Position and Velocity. Journal of Neurophysiology 91:515–532.

Perrault TJ, Vaughan JW, Stein BE, Wallace MT (2003) Neuron-Specific Response Characteristics Predict the Magnitude of Multisensory Integration. Journal of Neurophysiology 90:4022–4026.

Perrault TJ, Vaughan JW, Stein BE, Wallace MT (2005) Superior colliculus neurons use distinct operational modes in the integration of multisensory stimuli. Journal of Neurophysiology 93:2575–2586.

Pitkow X, Angelaki DE (2017) Inference in the Brain: Statistics Flowing in Redundant Population Codes. Neuron 94:943–953.

Pouget A, Snyder LH (2000) Computational approaches to sensorimotor transformations. Nature Neuroscience 2000 3:11 3:1192–1198.

Riedmiller M, Braun H (1993) Direct adaptive method for faster backpropagation learning: The RPROP algorithm. 1993 IEEE International Conference on Neural Networks pp. 586–591.

Salinas E, Abbott LF (1994) Vector reconstruction from firing rates. Journal of computational neuroscience 1:89–107.

Salinas E, Sejnowski TJ (2001) Gain modulation in the central nervous system: where behavior, neurophysiology, and computation meet. The Neuroscientist : a review journal bringing neurobiology, neurology and psychiatry 7:430–440.

Sanger TD (1996) Probability density estimation for the interpretation of neural population codes. Journal of neurophysiology 76:2790–2793.

Scott Murdison T, Leclercq G, Lefèvre P, Blohm G (2015) Computations underlying the visuomotor transformation for smooth pursuit eye movements. Journal of Neurophysiology 113:1377–1399.

Seung HS, Sompolinsky H (1993) Simple models for reading neuronal population codes. Proceedings of the National Academy of Sciences 90:10749–10753.

Snippe HP (1996) Parameter Extraction from Population Codes: A Critical Assessment. Neural Computation 8:511–529.

Stanford TR, Quessy S, Stein BE (2005) Evaluating the Operations Underlying Multisensory Integration in the Cat Superior Colliculus. Journal of Neuroscience 25:6499–6508.

Stein BE, Stanford TR (2008) Multisensory integration: Current issues from the perspective of the single neuron.

Trott AR, Born RT (2015) Input-Gain Control Produces Feature-Specific Surround Suppression. Journal of Neuroscience 35:4973–4982.

Ursino M, Cuppini C, Magosso E, Serino A, Pellegrino G (2009) Multisensory integration in the superior colliculus: a neural network model. Journal of computational neuroscience 26:55–73.

Wolpert DM, Ghahramani Z, Jordan MI (1995) An internal model for sensorimotor integration. Science (New York, N.Y.) 269:1880–1882.

Wu S, Nakahara H, Amari SI (2001) Population Coding with Correlation and an Unfaithful Model. Neural Computation 13:775–797.

Xing J, Andersen RA (2000) Models of the posterior parietal cortex which perform multimodal integration and represent space in several coordinate frames. Journal of cognitive neuroscience 12:601–614.

Yeh CI, Xing D, Williams PE, Shapley RM (2009) Stimulus ensemble and cortical layer determine V1 spatial receptive fields. Proceedings of the National Academy of Sciences of the United States of America 106:14652–14657.

Zemel RS, Dayan P, Pouget A (1998) Probabilistic Interpretation of Population Codes. Neural Computation 10:403–430.

Zipser D, Andersen RA (1988) A back-propagation programmed network that simulates response properties of a subset of posterior parietal neurons. Nature 331:679–684.

